# 3D printed SAXS chamber for controlled in-situ dialysis and optical characterization

**DOI:** 10.1101/2022.04.19.488724

**Authors:** Tamara Ehm, Julian Philipp, Martin Barkey, Martina Ober, Achim Theo Brinkop, David Simml, Miriam von Westphalen, Bert Nickel, Roy Beck, Joachim O. Rädler

## Abstract

3D printing changes the scope of how samples can be mounted for small angle X-ray scattering (SAXS). In this paper we present a 3D printed X-ray chamber, which allows for in-situ exchange of buffer and in-situ optical transmission spectroscopy. The chamber is made of cyclic olefin copolymers (COC), including COC X-ray windows providing ultra low SAXS background. The design integrates a membrane insert for in-situ dialysis of the 100 µl sample volume against a reservoir, which enables measurements of the same sample under multiple conditions using an in-house X-ray setup equipped with a 17.4 keV molybdenum source. We demonstrate the design’s capabilities by measuring reversible structural changes in lipid and polymer systems as a function of salt concentration and pH. In the same chambers optical light transmission spectroscopy was carried out measuring optical turbidity of the mesophases and local pH values using pH-responsive dyes. Microfluidic exchange and optical spectroscopy combined with in-situ X-ray scattering enables vast applications for the study of responsive materials.

## 1. Introduction

Small angle X-ray scattering (SAXS) is a powerful technique for studying soft matter and biological materials at the nanometer scale (Hura *et al*., 2013; Jacoby *et al*., 2015; Kornreich *et al*., 2015; Safinya *et al*., 2015; Kornreich *et al*., 2016; Kikhney & Svergun, 2015). Apart from the high resolution, the key advantage of SAXS is its ability to resolve structures in solution under physiological conditions (Kler *et al*., 2012; Mertens & Svergun, 2010; Chappa *et al*., 2021; Brennich *et al*., 2019). To this end fluid samples are traditionally filled in X-ray capillaries made of quartz glass. Samples are sealed and placed under equilibrated conditions in a high-flux X-ray beam at the synchrotron or in-house X-ray sources. Quartz capillaries suffer from curvature; therefore precise positioning is crucial for replicable results. Moreover, these studies require open systems e.g., in time-resolved stop-flow and flow through experiments, where repeated measurements are being conducted at different time delays in microfluidic channels (Ab’ecassis *et al*., 2007; Kihara, 1994; Krishnamoorthy *et al*., 2019; Kirby *et al*., 2013; Blanchet *et al*., 2015). Structural changes in scattering can also be measured by rinsing in buffer within the microfluidic device and thus gradually changing local concentration of e.g., salt or pH (Skou *et al*., 2014; Junius *et al*., 2020). A disadvantage of flow systems is that they cannot handle liquid crystalline or viscous mesophases, like lipid membrane systems or aggregated samples. They are useful, however, to study the dynamics of aggregation, which are typically carried out in microfluidic devices with merging channels that allow continuous mixing (Toft *et al*., 2008). In these devices it is key that a precisely positioned X-ray beam captures the temporal-spatial relationship of chemical and physical reactions (Denz *et al*., 2018).

Here we design and test an in-situ SAXS dialysis chamber which allows the exchange of buffer conditions in the X-ray sample holder. The design created from 3D printed COC plastic fits to commercial dialysis membrane inserts and enables in-situ SAXS experiments under varying buffer conditions, such as salinity, pH, or osmolyte concentrations. We demonstrate modes of operation using responsive lipid and peptide amphiphile mesophases. Buffer exchange is controlled by diffusion on time scales on the order of one hour allowing for repeated measurements of samples at different conditions on an in-house setup. In contrast to capillaries, our design has flat windows, facilitating positioning of the X-ray beam and reducing the sample-to-sample variation in sample thickness.

In-situ SAXS dialysis allows for X-ray experiments on the same sample under continuously changing conditions, and thereby will decrease sample to sample variations in a series of SAXS measurements. In addition, the COC chamber design is comparatively easy to be adapted and relatively cheap to produce. There are many examples where the relevant time scale of structural rearrangement is slow and hence can also be followed using in-house X-ray sources, where radiation damage is minute, rather than synchrotron beamlines (Scattarella *et al*., 2021; Jacoby *et al*., 2015; Schmelzer Jr *et al*., 2000). In these cases, in order to preserve precious samples, it is beneficial to only exchange buffer conditions via dialysis and to follow the structural dynamics in-situ on the same sample.

In recent years several studies presented microfluidic X-ray chambers made out of polymer materials, in particular the amorphous and optically transparent cyclic olefin copolymer (COC) (Schwemmer *et al*., 2016; Koester & Pfohl, 2012; Silva *et al*., 2015; Ghazal *et al*., 2016; Denz *et al*., 2018). This approach, using an X-ray transparent chip, allows for in-situ sample mixing during SAXS measurements or in-situ dilution of, for example, proteins with a variety of buffers. COC materials are also frequently used in crystallography screening by using 96 well plates made out of COC (Lee *et al*., 2013; Joseph *et al*., 2011) Traditionally, micro-fluidic X-ray chambers use the well-established window material Kapton, which has rather low optical transparency and needs adhesives for attachment to the chamber. COC devices have the advantage that they can be manufactured using 3D printing. In addition, foils from the same material can be used as X-ray window material. Annealing the foils to printed structures provides a simple, precise and reproducible production of sample chambers. Most importantly, COCs are well-suited for X-ray applications and optical measurements due to their high optical transparency and low background scattering at relevant photon energies as compared to PDMS and SiO (Guha *et al*., 2012). Previous studies (Denz *et al*., 2018) have benchmarked COC versus Kapton windows with gold colloid measurements at two different synchrotron beamlines. The background subtracted scattering data of the COC device and of the Kapton device agreed very well. So far, no COC chamber has been reported that allows for buffer exchange.

## 2. Chamber design

Our key design idea was to create a SAXS compatible sample chamber with in-situ dialysis capabilities. In order to avoid impractical handling with loose dialysis membranes, we used commercial dialysis inserts for Eppendorf cups (Slide-A-Lyzer™ MINI 0.1 mL, ThermoFisher). The SAXS chamber was built to create a small sample chamber in close proximity to the dialysis membrane. These dialysis inserts provide a 500 µL liquid reservoir, which can be extended via external syringe pumps. The dialysis inserts have a standardized size, are easily mountable, and come in various different pore sizes. For our purposes we used a molecular weight cutoff of 3.5 kDa. The surrounding sample chamber is first 3D printed with COC using an Ultimaker 3 and then sealed on both sides with transparent windows made of COC foil (Fig. 1). Because both the chamber and the windows are made of the same polymers, they can be annealed by simply heating both for a few seconds on a hotplate at 160°C. For our chambers we use 2.85 mm thick Creamelt™ COC filament (Herz GmbH) and 50 *µ*m thick TOPAS™ 8007F-04 COC foil (Microfluidic ChipShop).

**Fig. 1.**
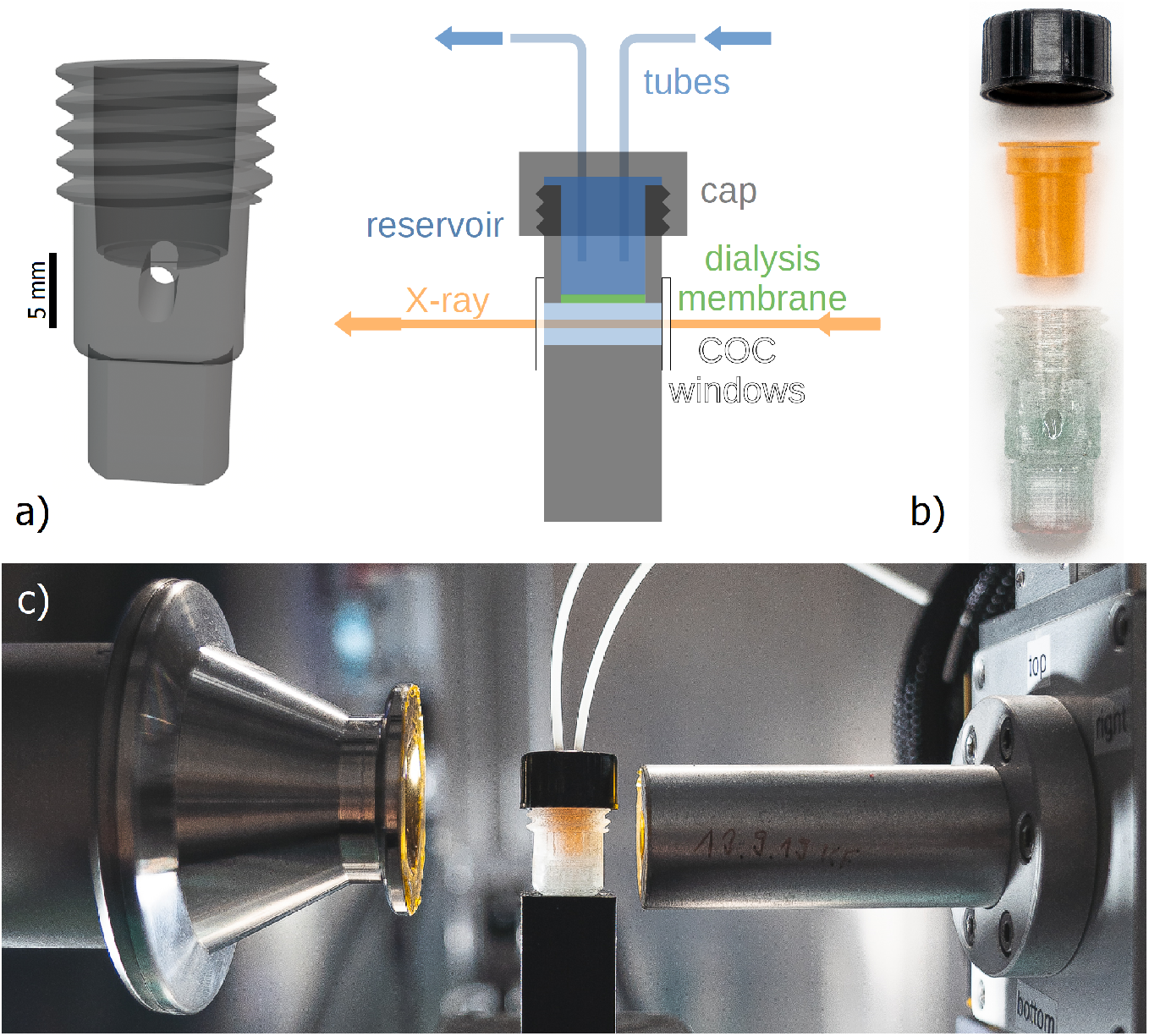
Chamber design principle and implementation. (a) Schematic drawing of the COC chamber. (b) Sample corpus is 3D printed out of COC to enclose the colored dialysis tube. Two COC X-ray windows allow for measurement of both condensed and and diluted phases. (c) Sample setup for in-situ measurements and dialysis using syringe pump systems.

The chamber can be filled from the top before closing it off by mounting the dialysis insert, which itself can be closed by a simple cap to avoid evaporation of the reservoir. The sample volume below the dialysis membrane is about 100 *µ*l and its optical path length is optimized for a 17.4 keV molybdenum source. If necessary, the reservoir can be continuously exchanged via syringe pumps and tubes through the cap to increase the reservoir volume. For time-resolved experiments, the chamber can be placed in the X-ray beam path where syringe pumps can exchange the reservoir buffer in-situ (Fig. 1).

We first characterized the signal-to-noise ratio of X-ray scattering in our dialysis SAXS chamber for both, COC or Kapton windows, using of small unilamellar vesicles made of SOPC at 30 mg/ml concentration (Fig. 2). The scattering signal is in agreement for both of the X-ray window materials, however, background measurements with the empty 3D printed chamber showed that Kapton windows have a slightly stronger background signal than COC. This is most visible in the *q*-regimes where Kapton is known to exhibit scattering, i.e. a broad peak around *q* = 0.09 Å^−1^ as well as in the WAXS region with scattering at *q* = 0.4 Å^−1^ depending of the type of Kapton.

**Fig. 2.**
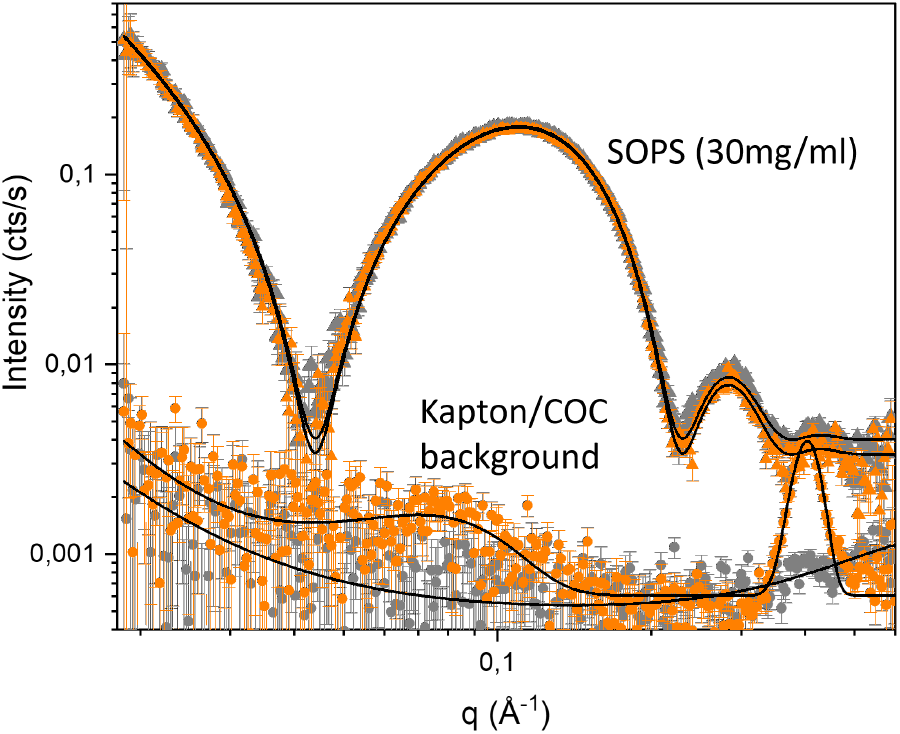
In-house measurements with Kapton and COC windows. Triangles: SAXS signal with errorbars (orange/grey: Kapton/COC windows) of SOPS LUVs with a diameter of 100 nm (30 mg/ml). Solid lines correspond to the best fit for flat symmetrical bilayers (see Supporting Information for model and detailed fit parameters). Circles: SAXS signal with errorbars of the empty dialysis chamber with 50 µm thick Kapton (orange) and 100 µm thick COC windows (grey). Solid lines correspond to the best fit for a power law with one and two Gaussians. The samples were measured with an exposure time of one minute per frame and measured for 6 hours in total.

### Buffer exchange kinetics

Buffer is exchanged between the reservoir and the sample volume via diffusive transport across the build-in dialysis membrane. For our design this reversible dialysis occurs over a time scale of a few hours. Our COC chamber allows for both optical and X-ray measurements of the sample in the same environment. We used this feature to follow the dialysis optically. Mixed at a ratio of 1:2, we used the pH indicators Chlorophenol Red and Bromothymol Blue, which change color from yellow to pink to blue for a pH range of 5 to 8. By preparing several samples within this range and measuring their transmission spectra in the chamber, we were able to calculate the spectra depending on the pH and vice versa (Fig. 3a). This allows for optical pH measurements without disturbing the sample during dialysis. Figure 3b shows the time evolution of pH inside our chamber during dialysis, which gives a good estimate of the diffusion times (*t*_1*/*2_ = 1 h). The result is in agreement with the calculated diffusion times 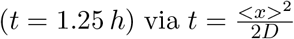, where *x* is the vertical diffusion path along the chamber (*x* ≈ 3*mm*) and *D* the typical diffusion coefficient for buffer salts 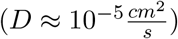.

**Fig. 3.**
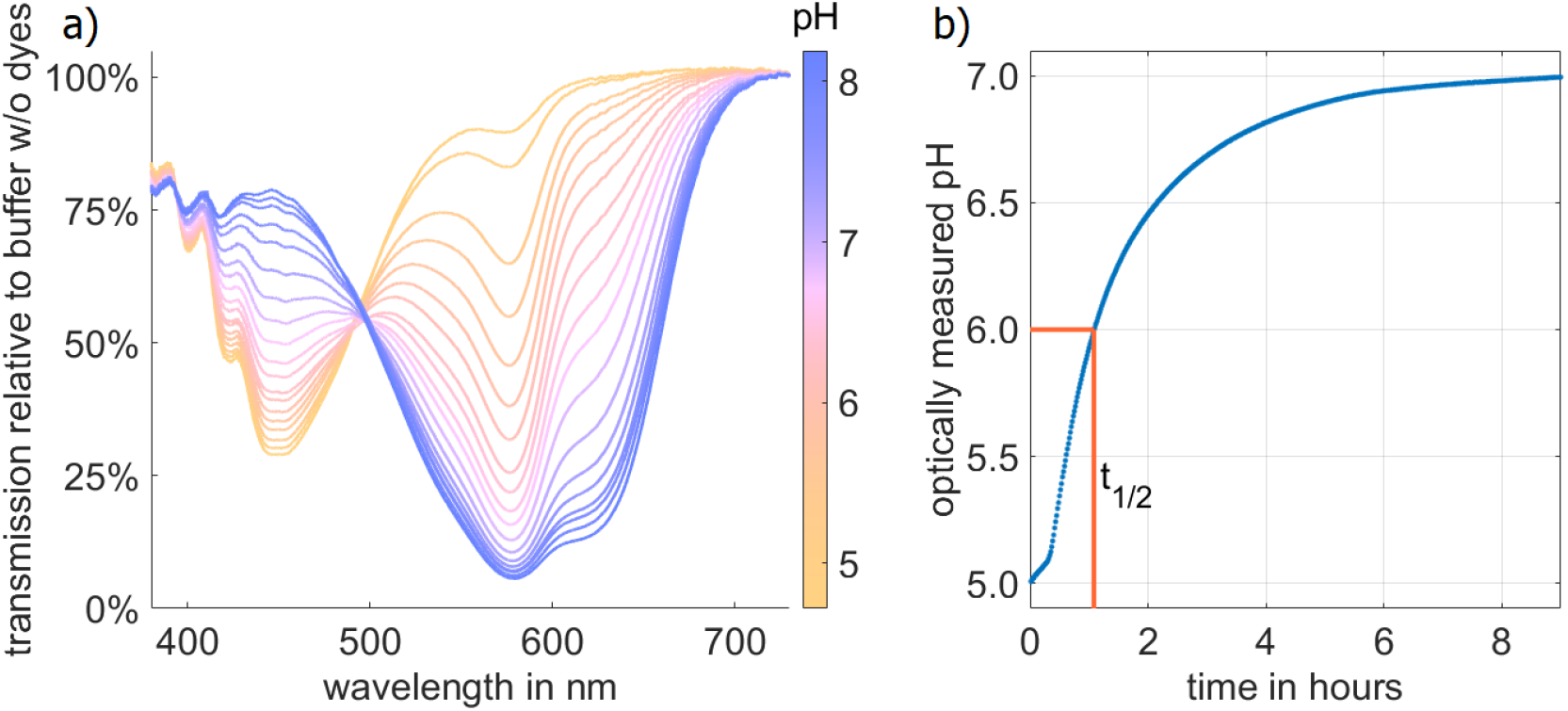
Time scale of in-situ pH change. a) pH-dependent transmission spectra of Bromothymol Blue and Chlorophynol Red mixed at a molar ratio of 2:1. b) Optical measurements of the pH inside the sample chamber during dialysis results in an approximate exchange halftime of *t*_1/2_ = 1 h.

To show reversibility, we used a pH-responsive system of intrinsically disordered peptide amphiphiles (IDPAs (Jacoby *et al*., 2021)). Specific amino acids in the used sequence can get protonated and become uncharged with decreasing pH. At pH 4.7 the IDPAs self-assemble into rod-like micelles. Upon increasing the pH to pH 7.5, the IDPAs transform into spherical micelles with a change in form factor scattering. We were able to monitor this transition from micellar rods to spherical rods and vice versa (pH 4.7 to pH 7.5) in an in-house SAXS source (see Supporting information). As the diffusion time scales for the transition points are larger than the time resolution of the in-house SAXS device, these experiments do not require a high resolution synchrotron beam. This makes measurement cycles easily applicable at home-sources. We monitored the transition with time-resolved SAXS measurements (Fig 4). The SAXS data didn’t show any changes after 6 hours.

**Fig. 4.**
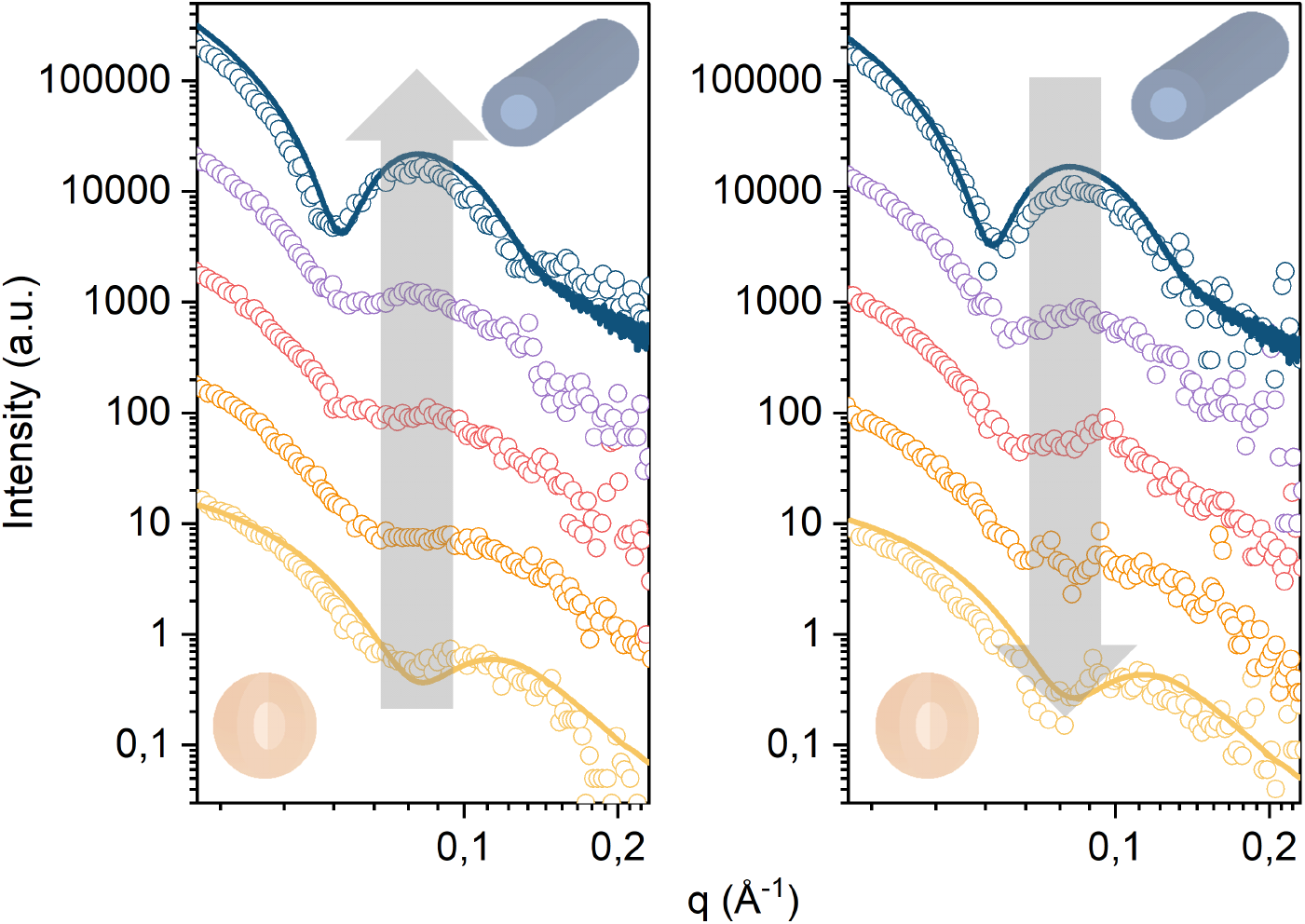
pH-dependent phase transition in an amphiphilic peptide mesophase. Left: Time-resolved in-house SAXS measurement (background corrected) of in-situ dialysis with 3D printed dialysis chamber from pH 7.5 to pH 4.7 of IDPAs shows a transition from micellar rods to spherical micelles in less than 6 hours. Scattering curves are snapshots at 0, 140, 160, 180 and 220 min with 20 min exposure time each, Solid lines are convoluted reference scattering curves from high resolution SAXS beamlines (Data was taken with an automated sample robot at DESY lightsource, Hamburg, Germany). Right: pH-dialysis back from pH 4.7 to pH 7.5 shows the reversibility of the transition in the dialysis chamber. Scattering curves are snapshots at 0, 260, 240, 340 and 520 min with 20 min exposure time each.

## 3. Experimental results

By conducting measurements with our 3D printed dialysis chamber, we demonstrated its usability in pH exchange. In principle, the chamber can be used for any buffer exchange where the diffusion molecules have a lower MW than the cut-off of the dialysis membrane, e.g. the exchange of salt often used in monitoring and controlling electrostatic interactions in biological SAXS experiments. With our 3D printed COC chamber, salt concentration series can now be performed in a single sample. An exemplary measurement is shown in Figure 5. Here, L-α-phosphatidylcholin (Soy PC) doped with 5 wt% of 1,2-Dioleoyl-3-trimethylammoniumpropan (DOTAP) was dialysed from 20 mM NaCl to 320 mM NaCl. The decrease in electrostatic repulsion due to charge screening resulted in shrinkage of the intermembrane distance (300 mM: 6.0 nm, 20 mM: 5.7 nm). The dialysis was reversed by exchanging the reservoir volume until the original salt concentration was restored. During this process, the lamellar repeat distance exhibited subsequent swelling.

**Fig. 5.**
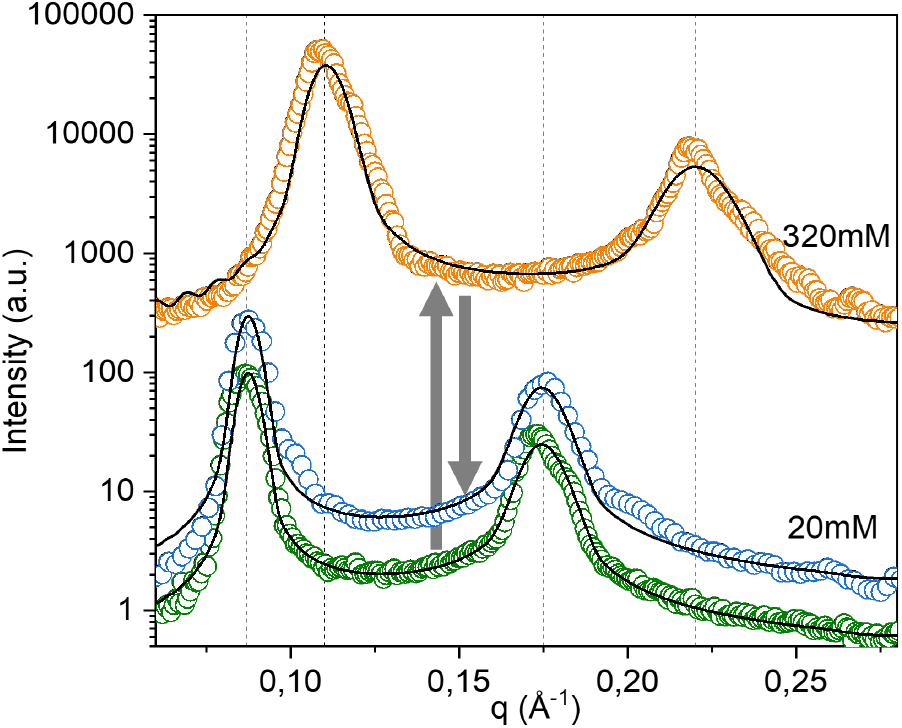
Reversible exchange of ionic buffer conditions. In-house SAXS SAXS signal of a charged lamellar phase (Soy PC doped with 5 wt% DOTAP) exhibits the shrinking and subsequent swelling of the intermembrane distance in response to increasing and decreasing salt concentrations. Black lines indicate fits of the data using the modified Caill’e theory (MCT) combined with a Gaussian electron density representation, as proposed in (Pabst *et al*., 2000). The Sample was equilibrated for 7 days.

Additionally, the design allows for absorbance measurements of (non) turbid samples. Thus time scales of phase transitions that exhibit a change in turbidity can be screened. Here, we examined the turbidity of the system of IDPAs presented in the last section. They undergo a phase transition from micellar, monodisperse, non turbid rods at intermediate pH into hexagonal packed rods at the isoelectric point (pH 4). The hexagonal phase has higher absorption rates, as these mesophases are in the size range of the wavelength of the applied light (Fig. 6).

**Fig. 6.**
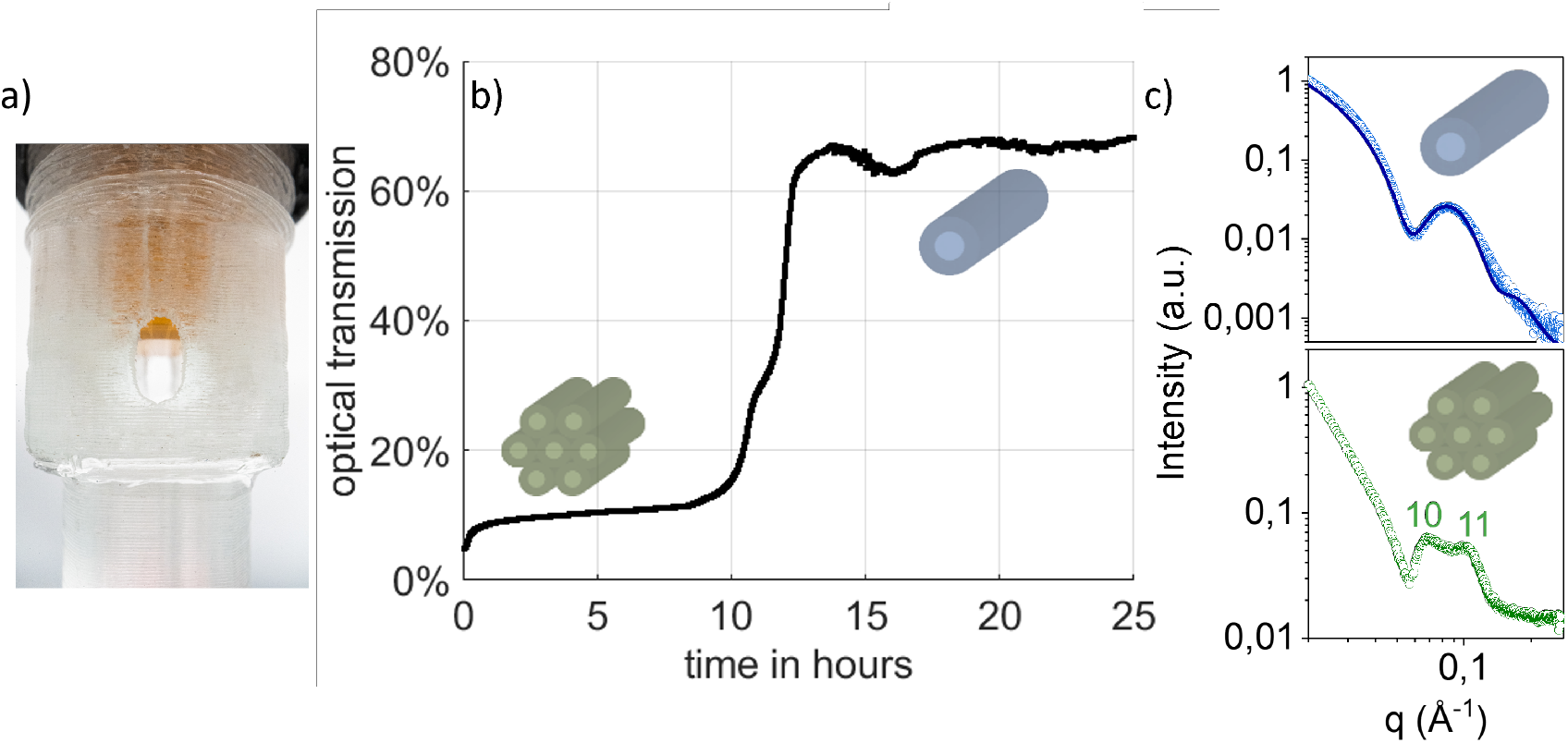
In-situ optical turbidity measurements. a) Photograph of the X-ray chamber showing the optical transparent X-ray window and the dialysis insert (orange). b) Optical turbidity across the X-ray chamber showing the structural transition from a condensed hexagonal to a dispersed cylindrical micelle phase. c) In-house SAXS signal of hexagonal-packed (green) and monodispersed micellar rods (blue) in solution min Data was taken with 20 min exposure time each. Solid line corresponds to spherical-core shell form factor (blue), Miller Indices show first second peak for hexagonal phase (repeat distance: 11.2nm)

**Fig. 7.**
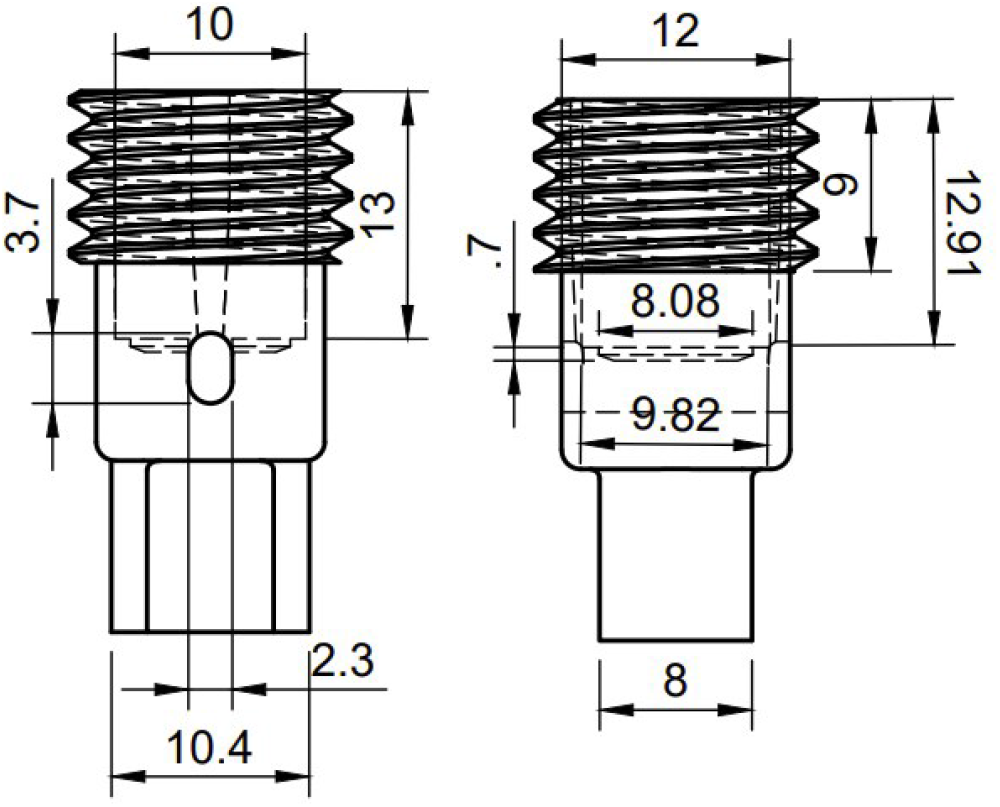
Technical drawing of 3D printed chamber. Front and side view of chamber, dimensions are given in mm.

## 4. Conclusion

In this paper, we designed and characterized a 3D-printed SAXS dialysis chamber made out of COC, and showed that the device is well-suited for in-house X-ray measurements. The chamber allowed for in-situ buffer exchange within a half-life time of about one hour as determined using pH-sensitive dyes. We showed that pH sensitive amphiphilic peptide mesophases underwent structural changes in a reversible fashion when pH was cycled from high to low pH and back to high pH. Likewise continuous change of ionic strength exhibited predicted shrinking and swelling of the spacing in charged lamellar lipid phases. Optical spectroscopy was carried out on the same samples within the X-ray chamber thanks to the optical transparency of the COC windows. The fabrication of larger numbers of X-ray chambers using 3D printing is simple and reliable. In contrast to closed capillaries, our 3D printed X-ray chamber is connected to a reservoir via a dialysis membrane and, therefore, allows for multiple measurements of the same sample under different buffer conditions. This approach saves sample material that is sometimes limited or costly. Most importantly using in-situ dialysis renders differential SAXS measurements highly reliable by avoiding sample-to-sample variations, which are typically caused by impreciseness in sample preparation or variations between sample holders. In our design the chamber dimensions were optimized for a Molybdenum 17.9 keV X-ray source with attenuation length in water of about 1cm (Bruetzel *et al*., 2016). In this case the integration of commercial dialysis insert was easily achievable. In principle a smaller sample volume is desirable since firstly, in-house SAXS sources with copper anodes could be used and secondly, diffusion limited buffer exchange would accelerate. However, a smaller sample volume requires smaller dialysis inserts and comes with the difficulties of laminating the chamber windows on an even smaller 3D-frame. Future chamber designs might overcome these problems. Alternatively, faster exchange kinetics in the existing design could be achieved by active mixing, e.g., using magnetic micro-stirrers. The later might also be required if dense samples exhibit strongly reduced diffusion times. In summary, 3D printed sample holders are promising for in-situ X-ray scattering providing microenvironments that enable the integration of continuous buffer exchange and inspection by optical spectroscopy.

## 5. Data availability statement

The cura profile to print is openly available in https://3dprint.nih.gov/discover/3dpx-016474, as version 01. Updates will be uploaded.

## 6. Supporting Information

The following parameters were used for printing the COC chamber:

|  |  |
| --- | --- |
| Printer | Ultimaker 3 |
| Material | COC 2.85 |
| Nozzle Size | 0.4 mm |
| Slicing | Cura 4.6.1 |
| Layer Height | 0.15 mm |
| Wall Thickness | 1.3 mm |
| Top/Bottom Thickness | 1.2 mm |
| Horizontal Expansion | -0.05 mm |
| Infill Density | 20 % |
| Printing Temperature | 245 °C |
| Build Plate Temperature | 80 °C |
| Printing Speed | 60 mm/s |
| Generate Support | None |
| Build Plate Adhesion Type | Brim |

### 6.1 Details about in-house Mo SAXS source

Our in-house SAXS system consists of a Mo GeniX3D microfocus X-ray tube (Xenocs SA, Sassenage, France) combined with FOX2D single reflection optics delivering a monochromatic and highly stable beam with an X-ray energy of 17.4*keV*. The flux is typically around 2.5 *×* 106 photons/s at the sample stage. For collimation the beam enters an 82*cm* long, fully evacuated collimation path closed by a 25 µm thick Kapton foil at the end. Collimation is achieved by integrating two partially motorized scatterless aperture slits (Xenocs SA, Sassenage, France), one upstream right at the mirror and the second at the tube exit.The sample stage is positioned 5 cm in front of the collimation path exit.

### 6.2 Small angle X-ray scattering (SAXS) analysis of SOPS LUVs data

All bilayer parameters are obtained from model fits of the background corrected total scattering intensity *I*(*q*) to an electron density profile Δ*ρ*(*z*) composed of three

Gaussians:

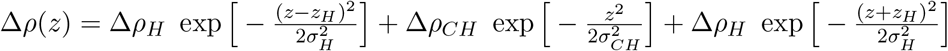

Here, Δ*ρ*_*H*_ is the scattering length contrast of the lipid head groups compared to water, i.e. Δ*ρ*(*z*) = Δ*ρ*_*H*_ − Δ*ρ*_*w*_. *z*_*H*_ is the spatial peak offset of the head centers in respect to the center of the bilayer, and *σ*_*H*_ is the corresponding variance of the Gaussian functions. Δ*ρ*_*CH*_ is the scattering length contrast of the lipid chains and *σ*_*CH*_ the variance of the Gaussian function describing the chain region. Model fitting was achieved by running the software-internal Levenberg-Marquardt algorithm using the software package SasView (http://www.sasview.org/).

**Table 2.** *Parameters obtained from least-squares fitting of SAXS data of 100 nm extruded SOPS vesicles to a symmetrical flat bilayer model*.

| Data | $z_H[\text{\AA}]$ | $D\rho_{CH}[a.u.]$ | $\sigma_{CH}[\text{\AA}]$ | $\Delta\rho_H[a.u.]$ | $\sigma_H[\text{\AA}]$ |
| --- | --- | --- | --- | --- | --- |
| COC | 21.1 | -0.002 | 8.6 | 0.004 | 4.3 |
| Kapton | 20.9 | -0.003 | 9.6 | 0.005 | 4.5 |

